# MSTD: an efficient method for detecting multi-scale topological domains from symmetric and asymmetric 3D genomic maps

**DOI:** 10.1101/370338

**Authors:** Yusen Ye, Lin Gao, Shihua Zhang

## Abstract

The chromosome conformation capture (3C) technique and its variants have been employed to reveal the existence of a hierarchy of structures in three-dimensional (3D) chromosomal architecture, including compartments, topologically associating domains (TADs), sub-TADs and chromatin loops. However, existing methods for domain detection were only designed based on symmetric Hi-C maps, ignoring long-range interaction structures between domains. To this end, we proposed a generic and efficient method to identify multi-scale topological domains (MSTD), including *cis-* and *trans-*interacting regions, from a variety of 3D genomic datasets. We first applied MSTD to detect promoter-anchored interaction domains (PADs) from promoter capture Hi-C datasets across 17 primary blood cell types. The boundaries of PADs are significantly enriched with one or the combination of multiple epigenetic factors. Moreover, PADs between functionally similar cell types are significantly conserved in terms of domain regions and expression states. Cell type-specific PADs involve in distinct cell type-specific activities and regulatory events by dynamic interactions within them. We also employed MSTD to define multi-scale domains from typical symmetric Hi-C datasets and illustrated its distinct superiority to the-state-of-art methods in terms of accuracy, flexibility and efficiency.

## Introduction

Folding of mammalian chromosomes into the nucleus has increasingly been recognized as an important factor in gene regulation, cell fate decisions, evolution and so on^1, 2, 3^. However, how chromosomes fold into the nucleus is still obscure. The chromosome conformation capture (3C) technique and its variants such as Hi-C, ChIA-PET and capture Hi-C have been employed to uncover the chromatin loops and hierarchical chromatin structural domains in three-dimensional (3D) genome architecture. Specifically, recent studies have revealed direct physical interactions, such as long-range chromatin contacts between enhancer and target genes^4^, actively co-regulated genes^5^ and Polycomb-repressed genes^6^. In addition, researchers have provided evidence for topologically associating domains (TADs) or sub-TADs^7, 8, 9, 10, 11, 12^. Furthermore, interactions between TADs at a variable distance result in active and inactive compartments, which are further subdivided into six different sub-compartments according to distinct patterns of histone modifications^4, 13^. Therefore, with the rapid accumulation of 3D genomic maps, developing efficient computational methods for detecting topological domains in chromosomal architecture is urgently needed.

Several methods have been developed to address these issues. For example, Dixon *et al.* adopted a hidden Markov model (HMM) method based on the directionality index from Hi-C maps to detect TADs^7^. Lévy-Leduc *et al.* proposed a block-wise segmentation model for TAD detection and proved that maximization of the likelihood on the block boundaries is a one-dimensional (1D) segmentation problem^14^. Similarly, Crane *et al.* developed an approach to transform the Hi-C contact matrix into an 1D insulation score vector for detecting topological structures^15^. Shin *et al.* employed an efficient and deterministic method to systematically identify TADs with a set of statistical measures to evaluate their quality^16^. However, these methods were only designed for detecting single-scale domains.

Recently, a few methods have been designed to explore the hierarchical organization of chromosomal architecture from symmetric Hi-C maps. For example, Filippova *et al.* introduced a dynamic programing model to identify hierarchical domains by adjusting a single parameter^17^. Weinreb *et al.* proposed a matrix decomposition model to infer a hierarchy of nested TADs based on ideal empirical distributions^18^. Zhan *et al.* presented a method CaTCH to detect hierarchical trees of chromosomal domains given a certain degree of reciprocal insulation from Hi-C maps^19^. However, these methods can only detect *cis*-interacting regions from Hi-C maps.

Very recent studies have revealed that both *cis-*and *trans-*interacting DNA regions exist in the cell nucleus^20^. With the emerging higher resolution genome-wide interaction maps of chromatin such as promoter capture Hi-C maps^21^ and ChIA-PET maps^22^, many distal DNA fragments regulate their targets bypassing long chromatin regions have been observed. Therefore, how to develop a generic method to infer both *cis*- and *trans*-interacting regions from diverse types of 3D genomic data is still a grand challenge in computational biology.

To this end, we proposed a generic and efficient method to identify **<u>m</u>**ulti-**<u>s</u>**cale **<u>t</u>**opological **<u>d</u>**omains (MSTD) from both asymmetric and symmetric 3D genomic datasets with a single adjustable parameter controlling domain scales. MSTD can detect long-range interacting domains such as those bewteen promoters and regulatory elements at variable distances, which can not been addressed directly by existing methods. We detected promoter-anchored interacting domains (PADs) from promoter capture Hi-C datasets across 17 primary blood cell types. We observed that the boundaries of these PADs can be well specified by one or the combination of a few epigenetic factors, indicating that these factors should play a key role the formation of genomic conformation. The analysis of the affinity relationship among cell types showed that PADs between functionally similar cell types have distinctly high conservation and consistent expression levels. Furthermore, dynamic PADs might perform specific cellular functions, while common ones take into account the underlying conditions of regular cell activities and participate in cell-specific regulatory events by dynamic interactions within each PAD. This suggests that PADs are important and basic units of genomic structure and function influencing gene regulation and cellular differentiation. We also employed MSTD to define multi-scale domains from symmetric Hi-C datasets with distinctly superior accuracy and efficiency compared to existing methods. Interestingly, the conservation of TADs between cells during continuous differentiation cycles is significantly higher than those between interval cycles. Last but not least, TADs are strongly correlated with epigenetic and transcriptional features.

## Results

### Illustration of MSTD

MSTD was inspired by a fast density-based clustering method, which is designed for grouping data points^23^. Here we group the strong contact interactions, which form square or rectangle submatrices corresponding to domains defined in Methods (Figure 1). Given a symmetric or asymmetric matrix, Firstly, for each diagonal element for a symmetric matrix in Figure 1C or a non-zero element for an asymmetric matrix in Figure 1E, MSTD computes two indexes: (1) local density defined as the average of the interaction frequencies with a cutoff radius window (Figures S1B, C and Supplementary Methods), and (2) minimum distance of the elements that have higher density (MDHD) than the element, Note that local density of the clustering center element is locally or globally maximum, and its MDHD are much larger than those of its neighbors. Thus, MSTD determines clustering centers with high MDHD by an adjustable parameter, which controls domain scales. Next, the remaining elements are assigned to the same cluster with their MDHD elements layer-by-layer. Meanwhile, individual outlier elements with high MDHD and low local density (Figure 1D) and the elements associated with these outlier elements will not be assigned to any cluster. Finally, the boundaries for each cluster are defined as the outermost elements of the same class in all directions.

**Figure 1.**
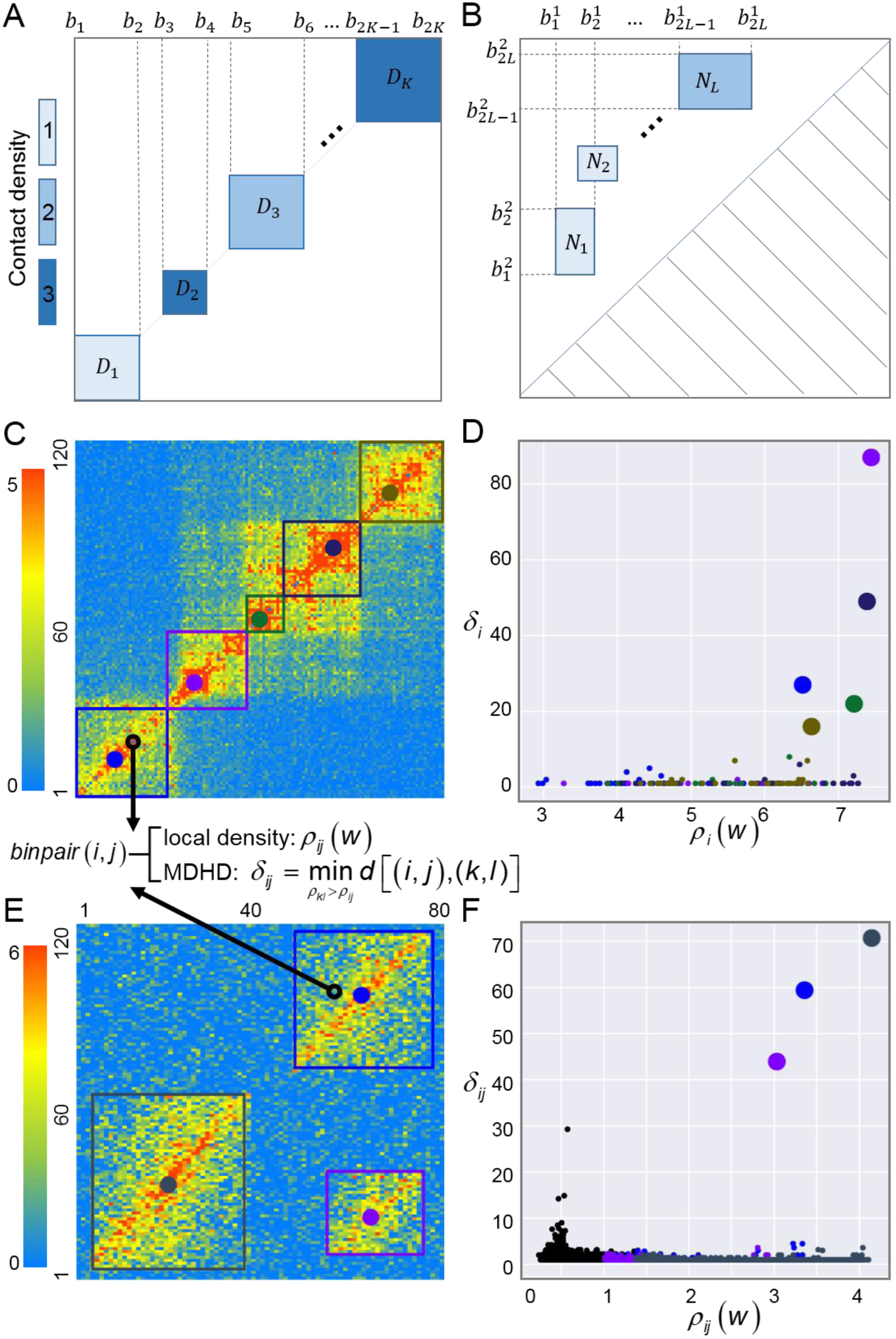
Illustration of MSTD for detecting TADs and PTADs in a symmetric and asymmetric matrix respectively. (A, B) Examples of TADs {*D*_1_,*D*_2_,⋯*D_K_*} and PTADs {*N*_1_,*N*_2_,⋯,*N_L_*}. The color depth represents the density of domains. (C) MSTD computes two indexes including local density and minimum distance of higher density (MDHD) for each diagonal element *k* in the symmetric matrix. (D) The decision graph of clustering results for (C). Clustering centers are marked by big dots with different colors. Starting with the centers, the remaining diagonal elements are assigned to the class of their MDHD element layer-by-layer. The clustering boundaries are defined to be the outermost elements of the same class in the two directions. (E) MSTD computes two indexes similar to (C) for each non-zero element *k* in the asymmetric matrix. (F) The decision graph of clustering results for (E). For the asymmetric matrix, MSTD employs all non-zero elements and defines the clustering boundaries in four directions. Noise elements were colored by black.

### Identifying PADs from asymmetric promoter capture Hi-C datasets

Firstly, we applied MSTD to identify multi-scale PADs from promoter capture Hi-C maps across 17 human primary blood cell types (Figure 2A and Supplementary Methods)^21^. MSTD can not only find *cis*-interacting PADs within the continuous genome, but also identify *trans*-interacting PADs between distal regulatory elements and target promoters (Figure 2A). The results below are based on parameter *δ_T_ =* 100 for detecting PADs, unless noted otherwise. MSTD costs only about 7.3 minutes for each chromosome (Table S1). Also, the size of the most of PADs detected by MSTD is in the range between 200kb and 2Mb (Figure S2A), as suggested by TADs from symmetric Hi-C maps in previous studies^8, 15^. Moreover, we noted that the same chromosome of different cell types obtains very similar number of PADs, especially between functionally similar cell types (Figure S2B), and average interaction frequency of intra-PADs is significantly higher than that of inter-PADs across 17 human primary blood cell types (Figure 2B), indicating that PADs relate to cell activities distinctly.

**Figure 2.**
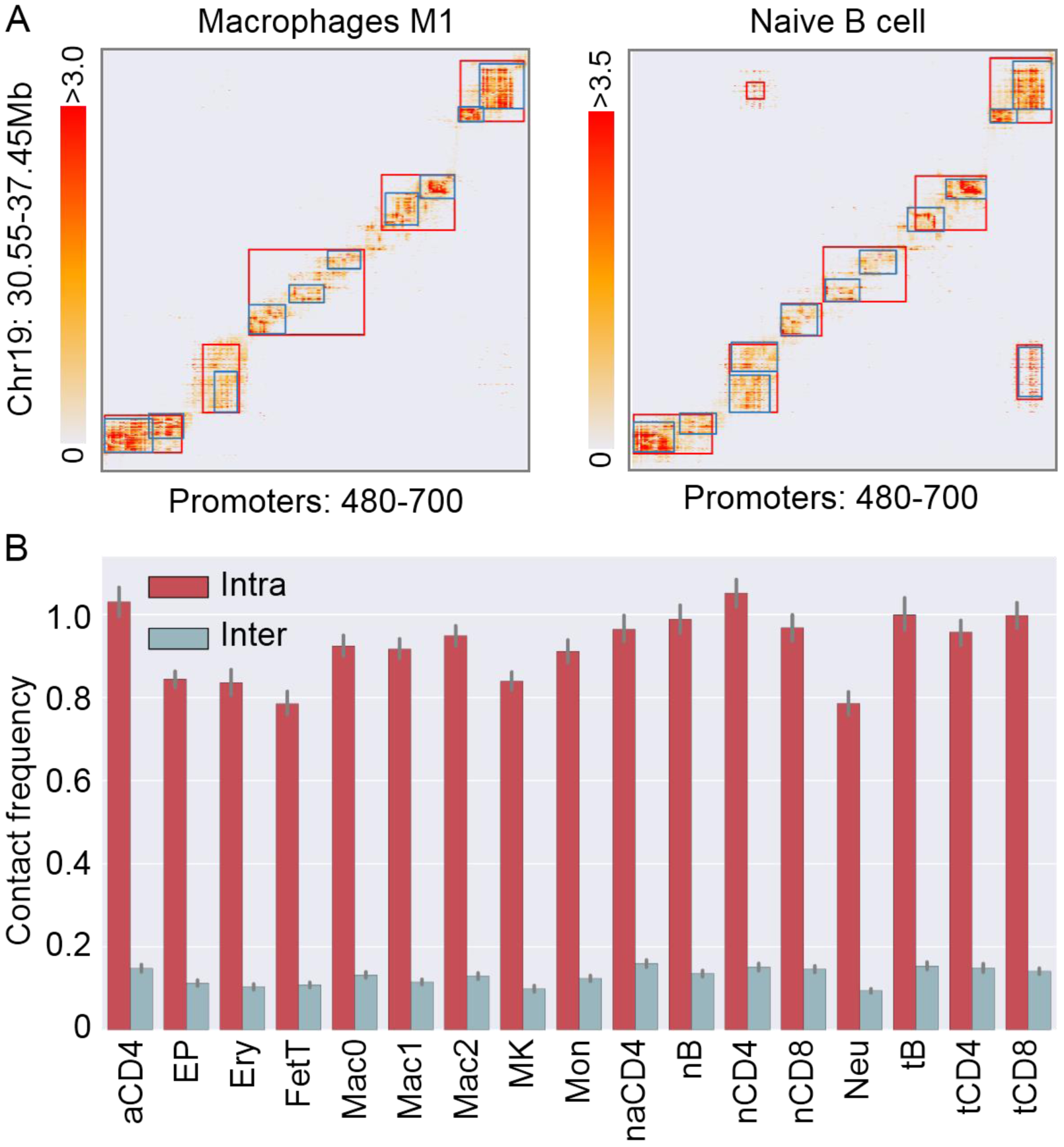
Illustration of PADs detected from promoter capture Hi-C datasets. (A) Examples of PADs detected by MSTD with *δ_T_* = 100 (blue) and *δ_T_* = 200 (red). (B) Comparison of the average contact frequency of *intra-*PADs and *inter-*PADs in 17 human primary blood cell types with *δ_T_* = 100.

### Epigenetic features is a good predictor of PADs

We defined the boundaries and centers of PADs along promoter interacting region (PIR) axis (Supplementary Methods) and found that epigenetic features show strong relevance to these PIRs’ regions compared with random ones (Figure 3, Figure S3-S5). Specifically, for monocytes cell type, the promoter mark H3K4me3 is strongly enriched in the boundaries of PADs (even more than twice that of intra-PADs) and the transcribed region mark (H3K36me3) is enriched in the regions slightly deviating from boundary regions, indicating that the boundary elements of PADs are significantly associated with promoters (Figure 3A). Meanwhile, the enhancer marks (H3K4me1, H3K27ac) are also enriched around topological boundaries, which might be related with chromatin loops in the boundary regions (Figure 3A). The transcribed region mark (H3K36me3) and the enhancer marks (H3K4me1, H3K27ac) demonstrate significant changes between intra-PADs and inter-PADs, suggesting PADs play key roles in active gene regulation and expression (Figure 3A). Meanwhile, the repressive mark (H3K27me3) is enriched in topological boundaries, which can be associated with the formation of repressive domains. Interestingly, the signals of H3K9me3 in inter-PADs are slightly higher than those in the intra-PADs, and both of which are significantly lower than random ones, revealing that PADs can segregate the spread of the heterochromatin (Figure 3A). Thus, multiple epigenetic features can identify different functional PAD boundaries. We also observed that multiple epigenetic features (except the heterochromatin mark) are enriched around the regions with high interaction intensity (PAD centers) (Figure 3A). Furthermore, we guess that one or the combination of several epigenetic features contributes to the formation of PADs, which can be associated with different functional units (Figure 3B). For example, H3K27ac tends to combine H3K4me3 or H3K4me1 to mark active boundaries of PADs, while H3K27me3 tends to mark repressive boundaries alone (Figure 3B). Therefore, histone modifications and DNase signal seem to be a good predictor of PADs and different functional units may have different strategies to specify chromatin domains.

**Figure 3.**
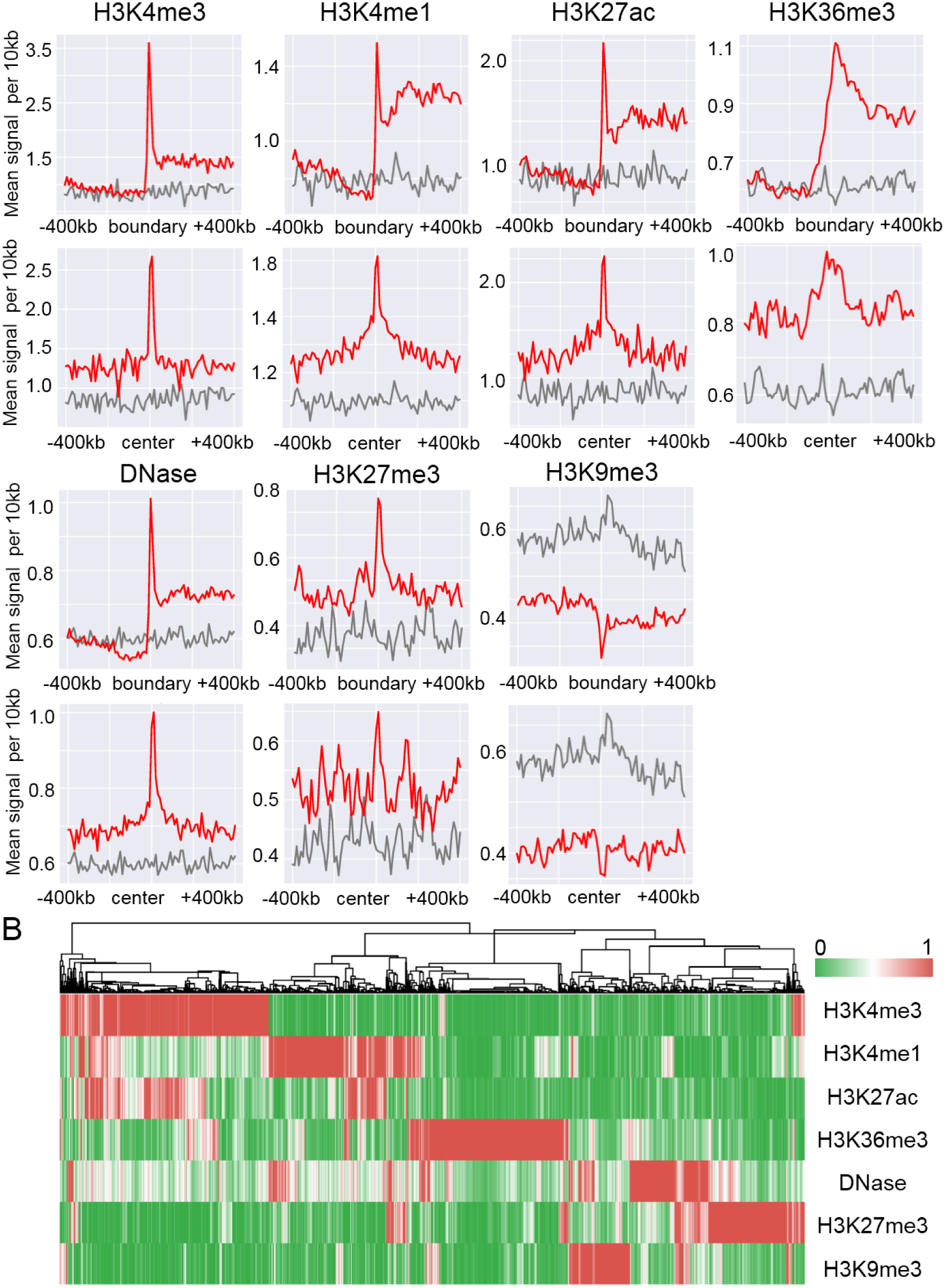
PADs are relating to epigenetic factors. (A) Enrichment analysis of seven epigenetic factors around boundaries and centers of PADs along PIRs in the human monocytes cell type. The observed and random signals are marked with red and gray colors, respectively. (B) Hierarchical clustering of boundaries of PADs according to their epigenetic factors.

### PADs can well capture the hematopoietic tree structure

The distribution of PADs among different cell types should reflect the affinity relationship of the 17 blood cell types. We regarded that the PADs are conserved between different cell types if their boundaries are adjacent (Supplementary Methods) and noted that 40%-70% of PADs are conserved between the same type of cells, yet only 30%-50% of PADs are conserved between the different type of cells, indicating the conservation and dynamics of chromatin structure during cell development and differentiation (Figure S6A). Furthermore, we derived five distinct classes based on the overlapping ratio of their PADs, which is highly consistent with the hematopoietic tree^24^, suggesting dynamic chromatin structures contribute to gene cellular functions (Figure 4A). The expression level of genes within these PADs distinguishes them into high (low) co-expressed ones and divided them into four categories of cell types consistent with hematopoietic tree, indicting these PADs play important roles in gene expression and regulation (Figures 4B, 4C and Methods)^24^. Interestingly, the gene sets tend to be very diverse across cell types within high and low co-expressed PADs respectively, indicting their underlying dynamical characteristics (Figure 4D). The above results reveal that the gradual change of the chromosomal structures (e.g., PADs) regulate the expression level of specific genes, which in turn promote the occurrence of cell development and differentiation. Therefore, it is important to distinguish these cell-specific PADs compared with stable ones for understanding the formation mechanism and cellular functions of PADs.

**Figure 4.**
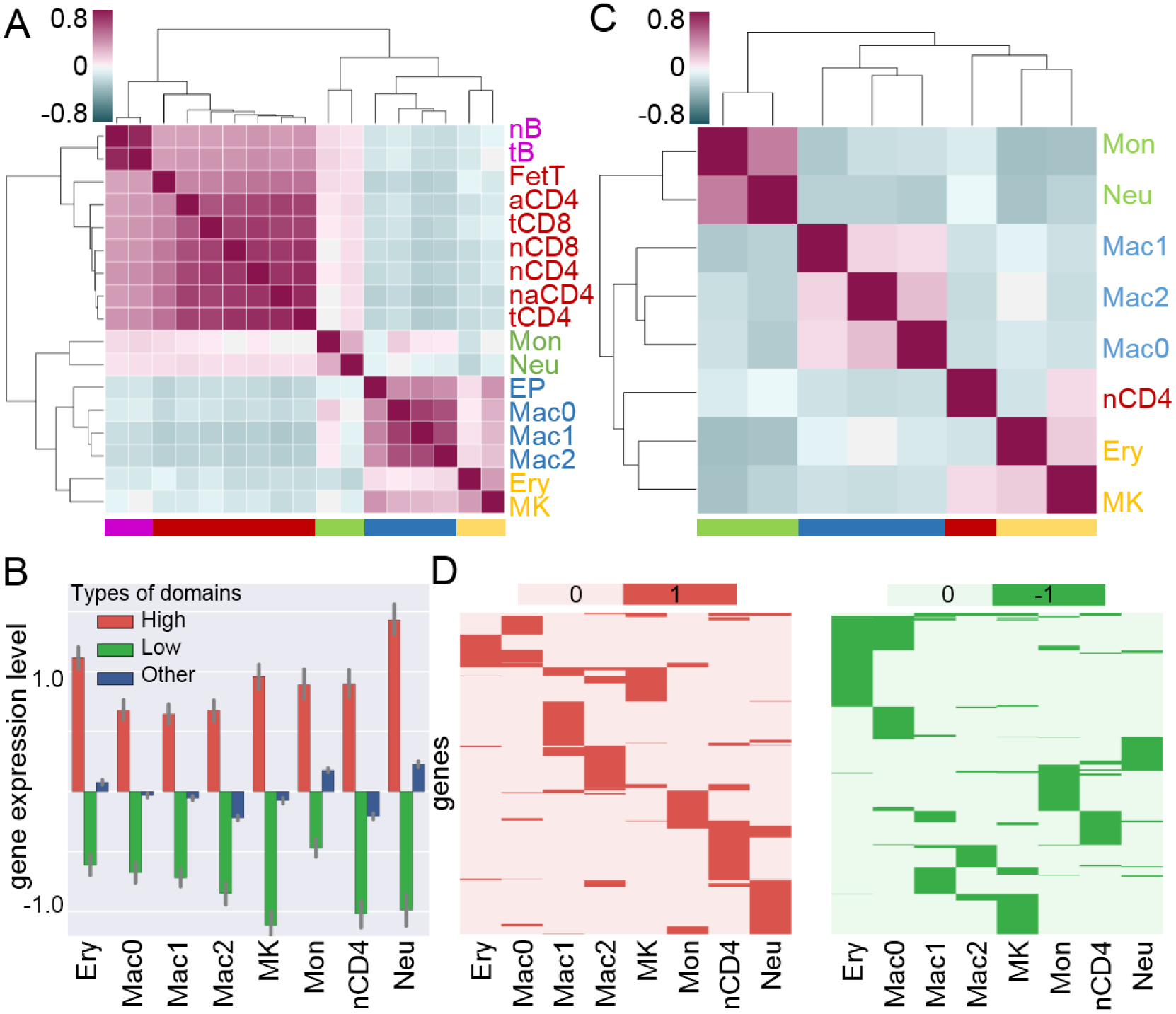
PADs can well capture the hematopoietic tree structure with distinct expression features. (A) Hierarchical clustering of the 17 cell types in terms of overlapping ratio of their PADs. Five clusters are marked by different colors. (B) The expression level of genes within high (low) expression level and other domains. (C) Hierarchical clustering of the eight cell types in terms of the correlation of z-score of PADs expression levels. (D) The genes of high (low) expression domains are marked by red and green respectively across eight cell types.

Next, we identified 693 lymphoid-specific, 426 myeloid-specific and 3808 common PADs according to the occurrence features among 17 human primary blood cell types to explore their cellular functions (Supplementary Methods). We further obtained 308 lymphoid-specific, 195 myeloid-specific and 1380 common PADs, whose pairwise chromatin regions containing more than one marker genes (Supplementary Methods and Table S2). Among those PADs, MSTD pioneered the discovery of 180 *trans-*PADs without any overlap between their pairwise chromatin regions (Figure 5, Figures S7-S14 and Tables S3 and S4).

**Figure 5.**
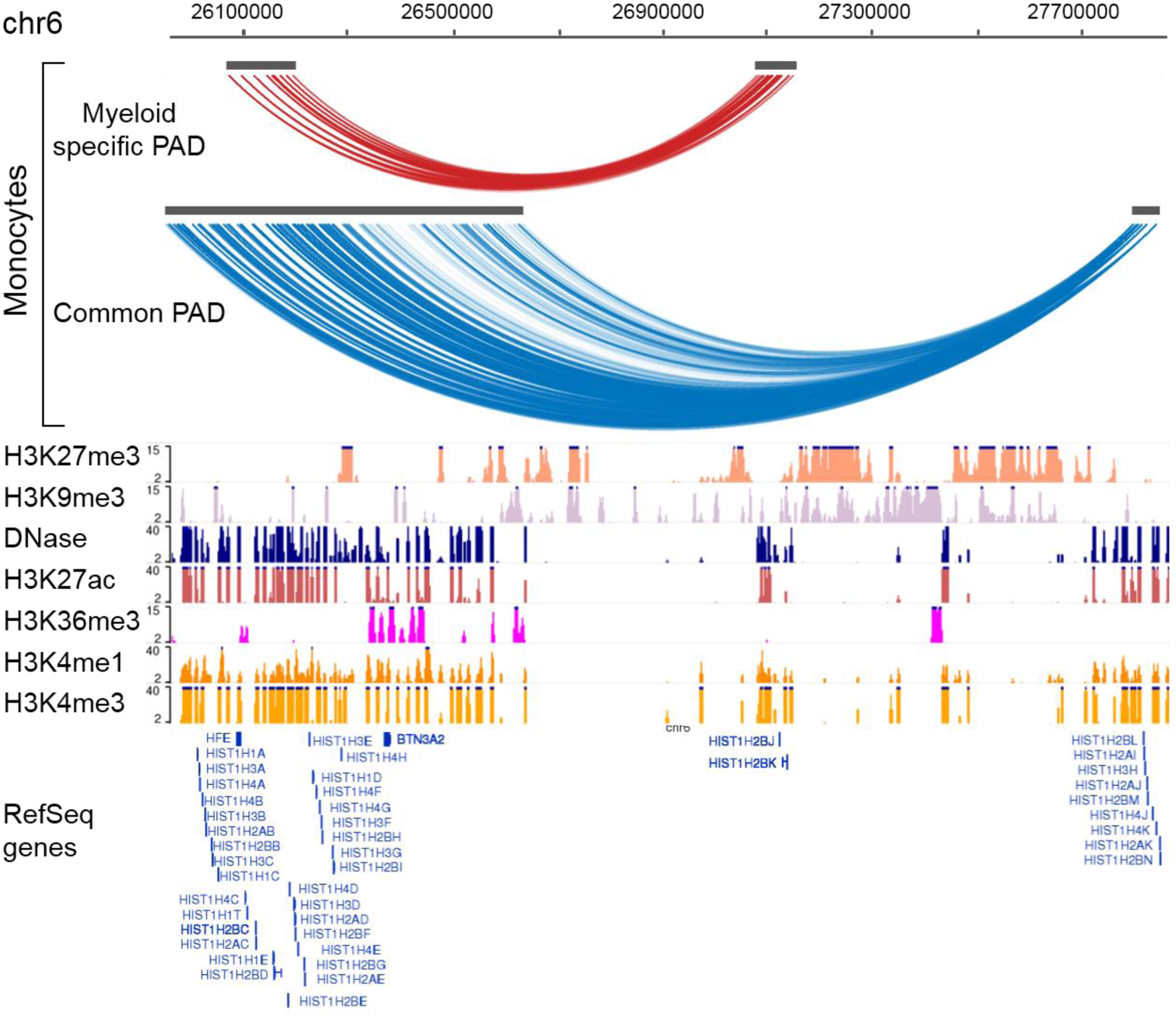
Illustration of a myeloid-specific PAD and a common PAD. Both PADs appears within the same genomic region using monocytes cell type as an example (chr6:26001891-27865483). Chromatin modification signals of seven epigenetic factors are shown in the monocytes cell type. The PADs connected with red arcs and blue arcs are a myeloid-specific one and a common one respectively. The image is drawn based on WashU epigenome browser with RefSeq gene annotations.

For example, we found two interesting *trans-*PADs within the same region of genome -- one myeloid-specific PAD and a common PAD shared by 16 cell types (except aCD4) (Figure 5 and Figures S7 and S8). For the myeloid-specific PAD, we can see that active TSS and transcription states occupies the pairwise chromatin regions of this PAD based on the ChromHMM annotations and active markers including H3K4me3, H3K27ac and DNase are significantly enriched in them too, which may suggest the chromatin loops within this PAD bring active promoters into the same nuclear space to initiate transcription^25^. Moreover, E2F, p53 motifs and E2F, E2F1 motifs are enriched in the upstream and downstream regions of this PAD, which contain genes related to innate immune response of myeloid tissues (upstream: HIST1H2BC, downstream: HIST1H2BJ and HIST1H2BK) (Table S3 and S4)^26^. This suggests that the chromatin loops among this myeloid-specific PAD play a special role in the cooperation between E2F1 and p53 to specifically activate innate immune response genes in myeloid tissues, which could promote the occurrence of apoptosis^27^.

For the lymphoid and myeloid common PAD, the pairwise chromatin regions contain several genes from histone families (H1, H2A, H2B, H3, and H4), which regulates circulating iron and mediates the regulation of lymphocyte and leukocyte (such as activation, cell-cell adhesion, immune response, antigen processing and presentation) (Figure 5 and Table S3)^26^. Furthermore, we identified 529 common and 40 dynamic chromatin loops between lymphoid and myeloid tissues from high-density chromatin loops of this PAD (*p*-value<0.01) (Supplementary Methods). Surprisingly, we found all of dynamic chromatin loops occur in lymphoid tissues and these dynamics interacting regions contains BTN3A2 gene involved in the adaptive immune response of lymphoid tissues (Figure S6B and Table S3)^26^. Thus, this PAD might participate in the underlying functions of regular activities of blood cells while enforce the adaptive immune response in lymphoid tissue. It can be seen that the upstream region of the common PAD contains that of the myeloid-specific PAD, and compared with the common PAD, the myeloid-specific PAD performs specific cellular functions by dynamic interactions with the different remote-region (Figure 5).

Another example is a lymphoid-specific PAD, which is characterized by a handful of active marks (Supplementary Methods and Figure S9). Both chromatin regions of this PAD contain more than one motifs associated with lymphocyte (Table S4). Specifically, GATA3 motif is a transcriptional activator, which binds to the enhancer of the T-cell receptor alpha and delta genes and Nur77 motif participate in modulating apoptosis in developing thymocytes^28^. Both GATA3 and Nur77 motifs are enriched in the upstream regions of this PAD, while TFE3 and SPIB motifs are enriched in its downstream regions. Previous studies have showed that TFE3 motif regulates T-cell-dependent antibody in aCD4 cell type and thymus-dependent humoral immunity, and SPIB motif is a lymphoid-specific enhancer^28^. In addition, Dontje *et al.* revealed that Delta-like1-induced Notch1 signaling pathway directs T cells and plasmacytoid dendritic cells decision by controlling the levels of GATA3 and SPIB^29^. More interestingly, within the genome region of the PAD, RC3H2 gene participates in T cell activation and differentiation involving in immune response and PTGS1 gene is related to blood pressure and circulation (Table S3)26. Thus, this PAD could perform lymphoid-specific functions by T cell specific marker gene expression, which are regulated by GATA3, SPIB and other transcription factors corporately (e.g. TEF3 and Nur77) (Supplementary Methods).

### Identifying multi-scale topological associated domains from symmetric Hi-C datasets

We next applied MSTD onto Hi-C datasets of mouse cortex cell binned at 40kb resolution^7^. MSTD can detect a few of larger topological domains with *δ_T_* increasing, indicating the existence of a hierarchy of structural domains (Figure 6A, Figures S15A and B). The average of contact frequency within a domain is significantly higher than that between domains and both of them decrease as the domain scale increasing (Figures S15C-E). Interestingly, inflection points appeared along parameter *δ_T_* between 5 and 10 for the two statistic measurements evaluating the quality of domains, which might be related to the presence of topologically associating domains (TADs) (Figures S15E and F).

**Figure 6.**
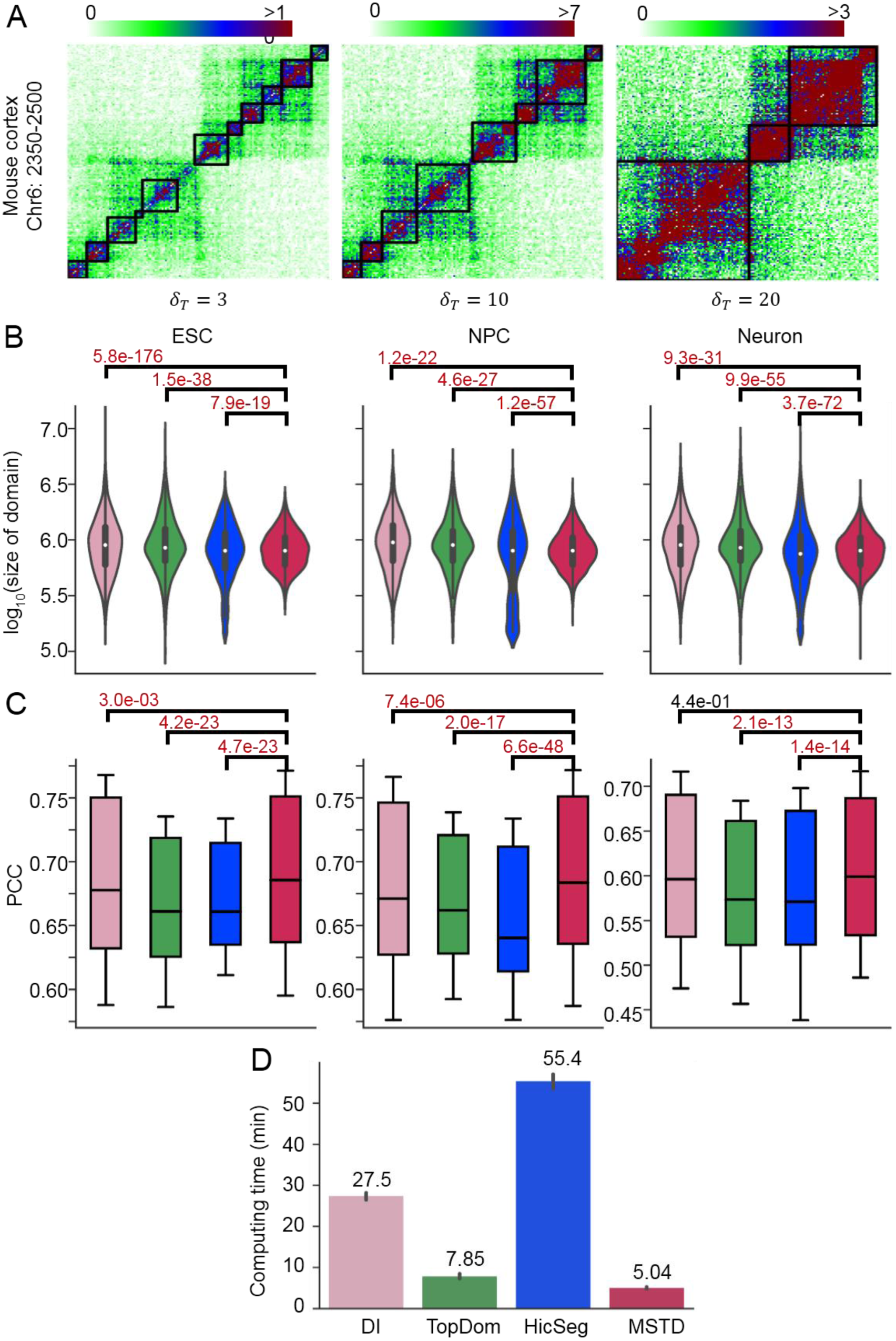
Illustration of multi-scale domains determined by MSTD from Hi-C datasets. (A) Multi-scale topological domain examples detected by MSTD with different MDHD thresholds (i.e., *δ_T_* = 3,10,20) of chr19: 2350-2500 bins of Hi-C maps in the mouse cortex cell (40kb resolution). (B) The size distribution of TADs detected by DI, TopDom, HicSeg, MSTD in three mouse cells (*p*-value<7.9 × 10^−19^, F-test). (C) Quality evaluation of TADs detected by the four methods in three cells using PCC measurements (*p*-value<3.0 × 10^−3^ except one case, T-test). The dark horizontal lines in boxes represent mean. The box surrounding each mean indicates the middle part of the data, which is the range from the 15th to 85th percentile. (D) Computing time of the four methods for domain detection across mouse ESC, NPC and Neurons cells.

### Comparison between MSTD and previous methods for identifying TADs

Previous studies suggested that the size of TADs should be between 200kb and 2Mb, which is used as one of the quality evaluation indicators^8, 15^. MSTD achieves remarkably higher consistency for all three cell lines compared with those of other methods with similar median of domain size at *δ_T_* = 7 (Table S5 and Methods). For example, we discovered 2965 qualified TADs, while DI, TopDom and HicSeq detect 1993, 1982, 2714 ones in Neuron cell line. Moreover, MSTD obtains TADs with smaller variance in domain scale (Figure 6B and Figure S16A).

Furthermore, MSTD shows superior performance than other three methods in terms of PCC and DIFF (Figure 6C, Figure S16D and Supplementary Methods), indicating that MSTD can detect more accurate TADs with coherent contact profiles. For a fair comparison, we employed the same number of domain boundaries of the top rank in terms of PCC and DIFF scores. MSTD generally shows much significant performance for all the three cell lines in the most cases (Figures S17 and S18). MSTD supposes that domain centers are surrounded by elements with lower local density and that they are relatively far away from the higher density elements. Thus, MSTD can well identify more consistent local density areas, which coincides well with compacted and elongated domain conformations in biology^23, 30^.

MSTD is friendly to users in parameter selections with very short computing time. Specifically, users can set an adjustable parameter *δ_T_* to control the domain scale (Figure S15 and Supplementary Methods). Meanwhile, MSTD takes about 5.04 minutes, while TopDom takes about 7.85 minutes, DI takes about 27.5 minutes and HicSeg spends 55.4 minutes approximately for detecting TADs of all chromosomes of three cell lines (mouse ESC, NPC and Neurons) in same computer environment (inter core 3.4GHz and 24G RAM) (Figure 6D).

### TADs show distinct biological conservation and specificity among ESC, NPC and Neurons

Previous studies have shown that TADs are conserved between cell types^7, 31^. We observed the average of common boundary ratio (CBR) is about 50%-80% and the average of TADs overlap ratio (TOR) is approximately 35%-70% during cell differentiation, which is similar with previous studies (Dixon et al., 2012) (Supplementary Methods and Figure S19). Interestingly, the conservation between cells during continuous differentiation cycle is significantly higher than those between the interval cycles. It will be useful to further investigate the result and extend it to specific functional and evolutionary events^32^.

### TADs are strongly correlated with epigenetic patterns and expressed features

The conservation of TADs during cell differentiation promotes us to explore the mechanisms of TADs formation. Recent studies showed that a common property of CTCF and highly active housekeeping genes may create insulating force of TAD boundaries^31, 33^. Intriguingly, architectural protein CTCF is indeed highly enriched at TAD boundaries in mouse ES and cortex cells (Figure 7A and Figure S20), and housekeeping exons and genes tend to be located in the boundary region of human ES cell (Figure 7B). Furthermore, H3K4me3, H3K9ac and RNA Polymerase II are significantly enriched and H3K36me3 is slightly enriched in the boundary regions of TADs for mouse ES and cortex cells, indicating the formation of TADs is indeed promoter-related^31^. Meanwhile, we found that transcription start sites (TSS) and expressed genes (FPKM>3) are also enriched around TAD boundaries (Figures 7A and 7B). Those observations reveal that factors associated active promoters and gene bodies can contribute to the formation of TADs boundaries (Figure 7A and Figure S20). Moreover, enhancer-marks (H3K4me1, H3K27ac and EP300) show a bimodal distribution around TAD boundaries (Figure S20). These peaks are located at 100kb-200kb away from the TADs boundaries, which may be associated with domains of gene co-regulation patterns^31^. H3K4me1 has a different bimodal patterns between mouse ES and cortex cells, which may be attributed to H3K4me1 marks sequences with region of DNA methylation loss in human mesenchymal stem cells (Figure S20)^34^. Interestingly, open chromatin mark (DNase signal) shows a significant enrichment in TAD boundaries of mouse ESC cell and a three peaks distribution around TADs boundaries of cortex cell (Figure 7A and Figure S20). In addition, inactive chromosome mark (H3K27me3) shows a slight peak in TADs boundaries for mouse ESC cell but not for cortex cell, which may be due to the responsibility of H3K27me3 for the repression of genes involved in cellular development and differentiation (Figure 7A and Figure S20)^35^. Finally, H3K9me3 is the mark of heterochromatin and shows no enrichment in both mouse ES and cortex cells (Figure 7A and Figure 20).

**Figure 7.**
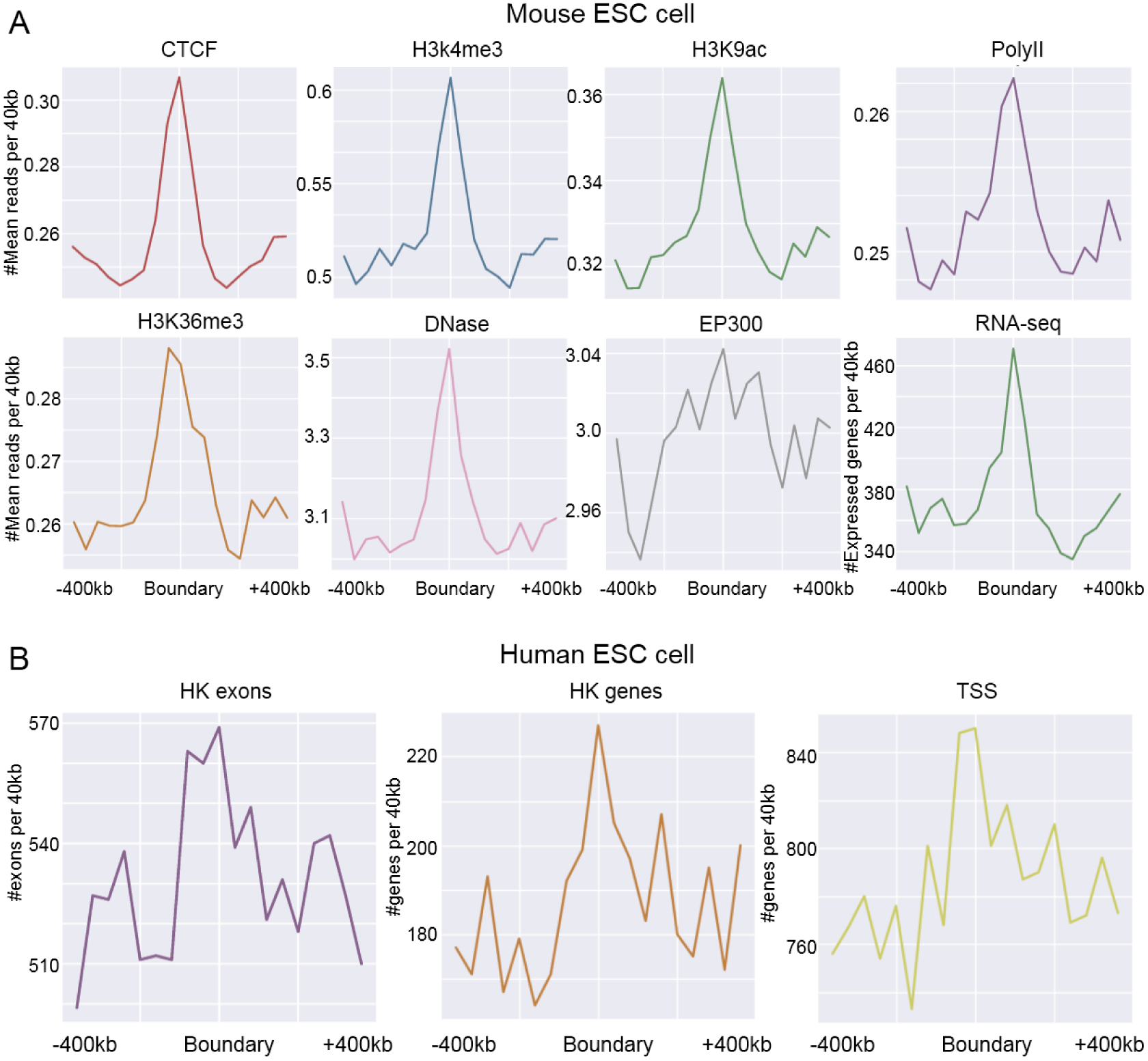
TADs are strongly related to epigenetic patterns and expression features. (A) The enrichment analysis of architectural protein CTCF, histone modifications, RNA Polymerase II, DNase-seq, EP300 elements and expressed genes around the boundaries of TADs identified by MSTD in mouse ES cell. (B) The enrichment analysis of housekeeping (HK) exons, HK genes and TSS of protein coding genes of hg19 RefSeq in GENCODE database in the boundaries of TADs identified by MSTD in human ES cell.

## Discussion

The chromosome conformation capture (3C) and its variants produce billions of read pairs that are used to draw genome-wide chromatin contact maps for multiple tissues and species. The accumulation of these data poses unprecedented challenges to computational biologists. Different tools have been developed to detect domains from symmetric Hi-C maps^36^. However, these methods pay no attention to interaction structures between domains (*trans-*domains) directly. MSTD begin to explore a variety of chromatin structures, including multi-scale TADs, PTADs and PADs from diverse chromatin contact maps, especially promoter capture Hi-C maps. Exploring a variety of chromatin structures will help to better understand the formation of spatial chromatin architecture and distinguish cellular function units. Meanwhile, 3D genomic maps of higher resolution will contribute to the presentation of landscape in the nucleus clearly and accurately.

We believe that the identification of multi-scale *cis-* and *trans-*domains will provide insights into local chromatin structure and promote the construction of 3D genomic model. Meanwhile, the analysis combining a variety of functional factors gradually explain cellular functions and biological processes of these structural units. With the development of new technologies, more diverse chromatin data will be generated for solving unknown biological problems. For examples, single cell Hi-C maps provide an opportunity to study the conservation and heterogeneity from cell to cell, revealing more granular cellular functions and biological processes in cell cycle^37, 38^. MSTD is a very flexible and powerful framework, which is expected to be applied to such new data in near future.

## Methods

### Materials

We first downloaded a comprehensive catalog of capture Hi-C datasets of 31253 annotated promoters and 230525 unique promoter-interacting regions (PIRs) in 17 human primary blood cell types^21^. The datasets detect 698187 high-confidence unique promoter interactions, of which 9.6% are promoter-promoter interactions and 90.4% are promoter-PIR interactions. The 17 cell types are from different nodes of the hematopoietic tree, which can be roughly divided into two categories, including 9 lymphoid cell types (Megakaryocytes (MK), Erythroblasts (Ery), Neutrophils (Neu), Monocytes (Mon), Endothelial precursors (Endp), Macrophages M0 (Mac0), Macrophages M1 (Mac1), Macrophages M2 (Mac2)) and 8 myeloid cell types (Naive B cells (nB), Total B cells (tB), Fetal thymus (FetT), Naive CD4+ T cells (nCD4), Total CD4+ T cells (tCD4), Non-activated total CD4+ T cells (naCD4), Activated total CD4+ T cells (aCD4), Naive CD8+ T cells (nCD8), Total CD8+ T cells (tCD8)). We collected available histone modification ChIP-seq datasets, including H3K4me3, H3K4me1, H3K27ac, H3K36me3, H3K9me3, H3K27me3, H3K9ac and DNase-seq datasets for seven cell types (Neu, Mon, FetT, nCD4. tCD4, nCD8, tCD8) from the Roadmap Epigenetics project (http://egg2.wustl.edu/roadmap/web_portal/). In addition, we collected available RNA-seq datasets for 8 cell types (MK, Ery, Neu, Mon, Mac0, Mac1, Mac2, nCD4) from^21^.

We also collected two sets of Hi-C maps. The first dataset includes mouse cortex, ES and human ES cell types^7^. For mouse cortex and ES cell types, we collected ChIP-seq datasets from the ENCODE project (http://www.genome.ucsc.edu/ENCODE/), including the architectural protein CTCF, seven histone modifications (H3K4me3, H3K9ac, H3K36me3, H3K4me1, H3K27ac, H3K27me3, H3K9me3), RNA Polymerase II, EP300, DNase-seq and RNA-seq data from^39^. For human ES cell type, we obtained human housekeeping genes from^33^ and TSS of protein coding genes of hg19 RefSeq in GENCODE database. The second dataset contains proliferating mouse embryonic stem cells (ESC), intermediate neuronal precursor cells (NPC) and post-mitotic neurons (Neurons) Hi-C datasets through different differentiation periods, binned at 50kb resolution from^32^.

### Definition of chromatin domains for Hi-C and promoter capture Hi-C datasets

Generally, Hi-C maps are summarized as a symmetric matrix *A* with bins of a fixed width (e.g. 40kb). Considering the symmetry of Hi-C matrix, in which *A_i,j_*(1 ≤ *i* ≤ *n*,1 ≤ *j* ≤ *n*) represents the interacting frequencies between bins *i* and *j*, we first define the diagonal blocks *D_k_* of high intensity with relative strong interactions as the topologically associating domains (TADs) (Figure 1A),

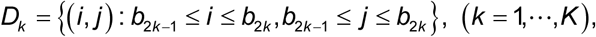

where *b*_2*k*–1_,*b*_2*k*_ (1 ≤ *b*_2*k*–1_ < *b*_2*k*_ ≤ *n*) represent the true TAD boundaries of the *k_th_* TAD and *K* is the number of TADs. We further define the non-diagonal blocks *N_l_* as pairwise topologically associating domains (PTADs) (Figure 1B),

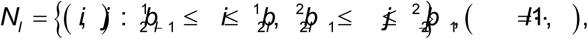

where 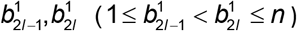 and 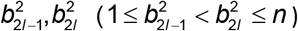 denote the boundaries of two distinct chromatin regions forming the *I_th_* PTAD and *L* is the number of PTADs. In this study, we denote promoter capture Hi-C map as an asymmetric rectangular contact matrix *B*, in which *B_i,j_* (1 ≤ *i* ≤ *n*,1 ≤ *j* ≤ *m*) represents interaction frequencies between promoters and PIRs. As a special type of PTADs, we define the special non-diagonal blocks *S_x_* as promoter-anchored interacting domains (PADs) from promoter capture Hi-C map (Figure S1A),

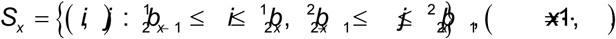

where 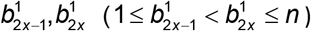 and 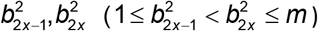 represent respectively the boundaries of promoters and PIRs forming the *x* th PAD and *X* is the number of PADs.

### MSTD

MSTD was inspired by a fast density-based clustering method, which is designed for grouping data points^23^. Here we group the strong contact interactions, which form square or rectangle submatrices corresponding to domains defined as above (Figure 1). Given a symmetric or asymmetric matrix *A*, where *A_i,j_* (1 ≤ *i* ≤ *n*,1 ≤ *j* ≤ *m*) denotes the interaction frequencies between bin *i* and *j*. Firstly, for each element (*i*, *j*) (a diagonal element (i.e., *i* = *j*) for a symmetric matrix in Figure 1C or a non-zero element for an asymmetric matrix in Figure 1E), MSTD computes two indexes: (1) local density *ρ_ij_* (*w*) defined as the average of the interaction frequencies with a cutoff radius window *w* (Figures S1B, C and Supplementary Methods), and (2) minimum distance *δ_ij_* of the elements that have higher density (MDHD) than the element (*i*, *j*),

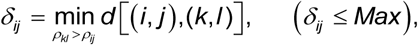

where *d*[(*i,j*),(*k,l*)] is the distance between the element (*i*, *j*) and element (*k*,*l*). We search for a MDHD element of the element (*i*, *j*) in the range of *Max*. Note that local density *ρ_ij_* of the clustering center element is locally or globally maximum, and its *ρ_ij_* are much larger than those of its neighbors. Thus, MSTD determines clustering centers by an adjustable parameter *ρ_T_* ( *δ_ij_* > *δ_T_*), which control domain scales. Next, the remaining elements are assigned to the same cluster with their MDHD elements layer-by-layer. Meanwhile, individual outlier elements with high *δ* and low *ρ* (Figure 1D) and the elements associated with these outlier elements will not be assigned to any cluster. Finally, the boundaries for each cluster are defined as the outermost elements of the same class in all directions. If the elements whose local density is significantly lower than their centers’, this element is removed from this cluster.

### Definition of high (low) co-expressed PADs

We first defined high (low) expression genes for a given cell type if their expression level is higher (lower) than their corresponding median of gene expression level across all available cell types. For each domain detected by MSTD, we defined a domain to be high- (low-) co-expressed domain by evaluating the difference between the number of high (low) expression genes in the genome region of this domain and the average number of high (low) expression genes in cyclically permutated genomes with the same number of genes as the domain region, which is weighted by its standard deviation (*p*-value<0.05).

### Parameter selection for MSTD and other methods

In order to compare MSTD with three popular methods including DI^7^, TopDom^16^ and HicSeg^14^ for identifying TADs, we next applied these four methods onto Hi-C datasets of mouse embryonic stem cells (ESC), intermediate neuronal precursor cells (NPC) and post-mitotic neurons (Neurons) through different differentiation periods (ESC-NPC-Neurons), binned at 50 kb resolution^32^. Since DI has been generally accepted to detect TADs in a number of existing studies, here we regard the size range of its TADs (from 700kb-1200kb) as the standard scale. In order to get a similar domain scale for a fair comparison, we set the adjustable parameters window=12 for TopDom, and *δ_T_* = 7 for MSTD, respectively. For HicSeg, we set the maximum number of boundary points (Maxk=chromosome length/850kb) and data distribution (distrib=’G’) respectively. Based on the above parameter settings, all four methods identify TADs with nearly identical domain scales (Table S5).

### Data and software availability

MSTD (version 0.0.2) is free, open source software under MIT License (OSI-compliant), is implemented in python3.6, and is freely available at https://github.com/zhanglabtools/MSTD or http://page.amss.ac.cn/shihua.zhang/software.html.

All datasets in this manuscript are public datasets. Their available link addresses are placed on Supplementary information.

## Acknowledgements

Yusen Ye would like to thank the support of the Academy of Mathematics and Systems Science at CAS during his visit. This work has been supported by the National Natural Science Foundation of China [No. 61532014, 61432010, 61672407 to LG; 11661141019, 61621003, 61422309, 61379092 to SZ]; the Strategic Priority Research Program of the Chinese Academy of Sciences (CAS) [No. XDB13040600], the Key Research Program of the Chinese Academy of Sciences [No. KFZD-SW-219], National Key Research and Development Program of China (2017YFC0908405) and CAS Frontier Science Research Key Project for Top Young Scientist [No. QYZDB-SSW-SYS008].

## Author contributions

SZ designed the study and proposed the main methods, YY performed the experiments, YY and SZ carried out bioinformatics and statistical analysis. YY, LG and SZ interpreted the results and drafted the manuscript. All authors read and approved the final manuscript.

## Additional information

**Supplementary Information** includes datasets sources, 20 figures, 5 tables and Supplementary Methods.

## Contact for reagent and resource sharing

Further information and requests for resources and reagents should be directed to and will be fulfilled by the Lead Contact, Shihua Zhang.

